# Relationship of sociodemographic and lifestyle factors and diet habits with metabolic syndrome (MetS) in a multi-ethnic Asian population

**DOI:** 10.1101/796144

**Authors:** Saleem Perwaiz Iqbal, Amutha Ramadas, Quek Kia Fatt, Ho Loon Shin, Wong Yin Onn, Khalid Abdul Kadir

**Author notes:** Corresponding author: Saleem Perwaiz Iqbal, Jeffrey Cheah School of Medicine and Health Sciences, Monash University, Malaysia.

## Abstract

**Objectives:** Literature shows a high prevalence of MetS among Malaysians, varying across major ethnicities. As sociodemographic characteristics, lifestyle factors and diet habits of such communities have been reported to be diverse, the study objective was to investigate the association of various sociodemographic characteristics, lifestyle factors and diet habits on MetS overall and among the three major ethnic communities in Malaysia.

**Materials and Methods:** We conducted a cross-sectional study among 481 Malaysians of age 18 years and above living in Johor, Malaysia. Information on demographics, lifestyle and diet habits was collected using a structured questionnaire. MetS was diagnosed among the subjects using the Harmonized criteria. Multiple logistic regression was used to analyse associations between sociodemographic and lifestyle factors and dietary behaviours with MetS.

**Results:** MetS was found among 32.2% of the respondents, more prevalent among the Indians (51.9%), followed by Malays (36.7%) and Chinese (20.2%). Overall, increasing age (AOR=2.44[95%CI=1.27-4.70] at 40-49 years vs. AOR=4.14[95%CI=1.97-8.69] at 60 years and above) and Indian ethnicity (AOR=1.95[95%CI=1.12-3.38)] increased, while higher education (AOR = 0.44[95%CI = 0.20-0.94] reduced the odds of MetS in this population. Quick finishing of meals (AOR=2.17[95%CI=1.02-4.60]) and low physical activity (AOR=4.76[95%CI=1.49-15.26]) was associated with an increased odds of MetS among the Malays and Chinese, respectively.

**Conclusion:** The population in Johor is diverse in these factors, and some of these are associated with MetS in certain ethnicities. In light of such differences, ethnic specific measures are needed to reduce the prevalence of MetS in this population.

## Introduction

Metabolic syndrome (MetS) is a combination of interrelated risk factors that predisposes individuals to the development of cardiovascular disease (CVD) and diabetes. This includes hyperglycemia, raised blood pressure, hypertriglyceridemia, low high-density lipoprotein (HDL)-cholesterol levels, and abdominal obesity, and is now recognized as a disease by the World Health Organization (WHO) and other international entities (1, 2).

According to a research across 7 European countries, the overall prevalence of MetS was estimated to be 23% using the WHO criteria (3). In Canada, nearly 25% of the adult population was found to be afflicted with MetS using the National Cholesterol Education Program (NCEP) – Adult Treatment Panel III (ATP III) (4). In Australia, the prevalence values of MetS using the WHO, NCEP-ATP III and International Diabetes Federation (IDF) criteria were 21.7%, 21.1% and 30.7%, respectively (5, 6). This points to the fact that prevalence of MetS within the same region may vary as different definitions are employed; and this could be due to the differences in the defined cut offs for its associated metabolic risk factors.

As the proportion and distribution of body fat in Asians, in general, were found to be different from Caucasians in North America and Europe, it became apparent that the definition of obesity applied to Western populations would not be applicable to Asian populations (7, 8). Therefore, the estimated prevalence values of MetS among Asians were found to be increased when Asian-adapted definitions of obesity were employed in the NCEP-ATP III. For example, in the Southeast Asian region, it increased from 13.1% to 20.9% for Singaporean males and for the Chinese adults, it increased from 10.1% to 26.3% (9, 10). A similar trend was observed among the Malaysians where during the 2008 nationwide survey an overall prevalence of 42.5% from 4341 subjects was reported using the Joint Interim Statement (JIS) “Harmonized” criteria, compared to 34.3% via the NCEP-ATP III criteria (11). Like in NCEP-ATP III, MetS according to the Harmonized definition includes any 3 of the 5 metabolic abnormalities – central obesity, hypertriglyceridemia, low HDL-cholesterol, high blood pressure and hyperglycemia (2). However, the Harmonized criteria have defined Asian cut-offs for central obesity (waist circumference: ≥ 90 cm for males; ≥ 80 cm for females) and reduced cut-off for hyperglycemia (≥ 5.6 mmol/L, instead of 6.1 mmol/L in the NCEP-ATP III). Ramli et al., using the Harmonized definition reported the prevalence of MetS to be 43.4% in 2013, from among 8,836 subjects across East and West Malaysia (12). This percentage was very close to the 42.5% prevalence reported in the 2008 nationwide survey (11).

The prevalence of MetS is dependent on a variety of non-modifiable (gender, age, ethnicity) and modifiable (lifestyle, diet) risk factors. These factors are known to, directly or indirectly, influence MetS among populations. For instance, Wen and colleagues reported the prevalence of Mets in rural China as 44.3% (by modified NCEP-ATP III criteria), 40.7% (by IDF criteria) and 47.7% (by Harmonized criteria), amongst a large cohort of 4748 subjects, primarily among females aged 50 years and above (13). From a study in Canada, Liu and colleagues reported MetS prevalence to be higher among Cree Indians compared to other aboriginal and non-aboriginal Canadians (14). Malaysia is no different as according to the nationwide survey the prevalence was found to be higher among older age groups, more among females, and most common among the Indians compared to other races in Malaysia (11).

Studies have shown that various lifestyle factors influence MetS. Sedentary lifestyle and physical inactivity have been shown to contribute to the development of MetS and its components (15–19). Smoking and alcohol consumption have also shown to have variable influences on MetS and its components (20–24). Furthermore, diet habits such as speed of eating, dining out, skipping breakfast and late dinners have been found to be associated with increased incidence of MetS (25, 26). These factors are present in most communities and might provide some insight on how their influence on MetS may be regulated in populations to contain its life-threatening complications. Reports mentioned above indicate the important influence of lifestyle habits on prevalence of MetS in a particular population. Malaysia is a unique country in Southeast Asia because of its ethnic diversity, culture, lifestyle choices and dietary intake habits. The influence of differing lifestyle choices and diet habits across the 3 major races of the country may provide a better understanding of the high prevalence of MetS in the country, along with measures for its containment. There have been only a few studies carried out in Malaysia on investigating the influence of lifestyle factors with the risk of MetS among the Malaysian population (27–29). While the two studies by Chu and Moy described the influence of physical activity on MetS on the Malays, the only study which dealt with ethnic differences with respect to physical activity and prevalence of MetS among the Malaysian population was based on the data that were collected more than 13 years ago (27–29). Moreover, in that study the relationship of lifestyle behaviours with MetS among major ethnicities was not reported (29). Therefore, the objective of the present study, therefore was to determine the association of sociodemographic characteristics, lifestyle factors and diet habits with the risk of MetS, overall and among the 3 major ethnic groups residing in Johor, Malaysia.

## Materials and methods

### Study design and location

This was a cross-sectional study, employing a nonprobability sampling strategy, conducted in Kulai and Felda Taib Andak of the Kulai district and Johor Bahru, Ulu Tiram and Kota Masai of the Johor Bahru district of Malaysia.

### Recruitment and eligibility criteria

Research camps were set up in central locations of Kulai and Johor Bahru districts, which were easily accessible to the target community. The selection of these study locations was partly based on the available percentages of MetS across each major ethnicity of Malaysia, reported in the nationwide survey 2008, to have enough subjects to represent each ethnicity in Johor so that data could be available for in-depth analysis for the stated objectives. Assistance was sought from community elders for making the locals aware of research camps and to convey the requests for their participation.

The inclusion criteria for the study were that the subjects should be of age 18 years or above, of either sex, and had been residing in Johor for at least one year. The subjects were requested to observe a 10 to 12-hour fast before arriving at the medical camp to donate blood samples for accurate assessment of fasting serum levels of glucose, triglycerides and HDL-cholesterol. Exclusion criteria included pregnancy or having any illness which could preclude their participation in the study such as cancer, liver disease, etc. Consented participants were invited to visit these camps for a physical examination and collection of blood samples upon observing a 10 to 12-hour fast. Upon sample collection, the subjects were asked about their lifestyle and dietary habits. Participants not observing the 10-12 hour fast were excluded from the analysis.

### Data collection and measurement

Data from the participants were collected using a structured questionnaire and proforma which contained information on anthropometric measurements, measurement of blood pressure, blood sample analysis results and questions on sociodemographic and lifestyle behaviors. The questionnaire was designed in English and back translated in the Bahasa Malaysia language. In the study, Bahasa language based version was used.

Body height was measured using Seca stadiometers (Seca, USA), while the weight was measured using the InBody 120 body fat analyzer (Biospace, Korea). Measures were taken to ensure that the subjects wore light clothing and no shoes. The measurement was recorded to the nearest 0.1 cm and 0.1 kg, respectively.

Waist circumference was measured using a measuring tape. The measurements were taken at the mid-point, between the lower rib margin (12^th^ rib) and the iliac crest. Caution was maintained during measurements that the subject was standing straight with feet together and arms relaxed on either side. It was also ensured that the tape was held in a horizontal position, wrapped around the waist, loose enough for the assessor to insert his/her finger between the tape and the subject’s body. The subject was instructed to breathe normally during the assessment, with the measurement recorded at the end of a normal exhalation and rounded to the nearest 0.1 cm.

Blood pressure was recorded using the Omron digital sphygmomanometers (HEM-7121, Omron Healthcare, Japan). The subject was provided a 4-5 minute rest, in a seated position, with the arm supported at heart level. At least two readings were taken from each subject, recording the concurrent or highest measurement obtained from the two readings. A third reading was taken in case, the difference between the two readings for systolic blood pressure was more than 10 mmHg, and for diastolic blood pressure more than 5 mmHg.

Fasting blood samples were collected from study participants for determining the levels of fasting serum glucose (in mmol/L), fasting serum triglycerides (in mmol/L) and fasting serum HDL-cholesterol (in mmol/L). Standard guidelines for phlebotomy were followed throughout the venipuncture procedure (30).

Collected samples were transported in cold chain to the laboratory where these were centrifuged, and the sera samples were separated and placed in identity marked cryotubes or Eppendorf tubes. These were then placed in a −60 degree Celsius freezer till laboratory analysis.

The blood analysis for determination of serum levels of fasting glucose (mmol/L), triglycerides (mmol/L) and HDL-cholesterol (mmol/L) was carried out using clinical chemistry analyser (Cobas C III). Its reagents were purchased from Randox Laboratories, United Kingdom.

The participants’ physical activity status was inquired using the International Physical Activity Questionnaire (IPAQ) (31). The questionnaire contained seven questions; the first two pertaining to the time spent on vigorous activities performed, the next two for moderate activities, the next two for mild activities and the last question was on the time spent while sitting. Responses were converted to Metabolic Equivalent Task minutes per week (MET-min/week) according to the IPAQ scoring protocol. The protocol also provides details for data processing, cleaning and truncation. Total minutes over last seven days spent on vigorous, moderate, and mild activities were multiplied by 8.0, 4.0, and 3.3, respectively, to create MET scores for each activity level. MET scores across the three sub-components were then summed to indicate the overall physical activity score. These overall scores were then categorized into high (total activity of at least 3000 MET-min/week), moderate (total activity of at least 600 MET-min/week) and low (total activity < 600 MET-min/week).

Diet habits included quick finishing of their meals, frequency of late dining, frequency of skipping breakfast and frequency of dining out. For quick finishing of meals, the question was asked on the subject’s perception on finishing their meals either fast (less than 10-15 minutes) or not fast (32–34). The assessment of the other three diet habit questions (frequency of late dining, frequency of skipping breakfast and frequency of dining out) were based on the participants’ frequency per week; three times or less were considered favorable (32). “Late dining” was defined as a meal eaten within two hours before bed-time. “Dining out” was defined as a meal consumed by the participant that is not prepared at his/her home (35–37).

### Definition of MetS

MetS was defined using the Harmonized criteria as having at least 3 of the following 5 risk factors: 1) Abdominal obesity, defined as having a waist circumference ≥ 90 cm for males and ≥ 80 cm for females; 2) Raised serum triglycerides (hypertriglyceridemia), defined as 1.7 mmol/L (150 mg/ dL); 3) Low high density lipoprotein cholesterol (HDL-C), defined as 1.0 mmol/L (40 mg/dL) for males and 1.3 mmol/L (50 mg/dL) for females; 4) Raised blood pressure, defined as a systolic blood pressure ≥ 130 or a diastolic blood pressure ≥ 85 mmHg, or current use of anti-hypertensive medications; and 5) Raised fasting blood sugar (hyperglycemia), defined as ≥ 5.6 mmol/L (100 mg/ dL) or current use of anti-diabetic medications.

### Statistical analysis

Data entry was performed using EpiData version 3.1. During the process of data entry, 5% of the forms were re-checked for accounting any errors during entry of data. All data were analyzed using Statistical Package for Social Sciences (SPSS) version 23 (IBM SPSS Statistics for Windows, Version 23.0. Armonk, NY: IBM Corp.).

The sample size estimate was calculated using estimates of various components of MetS reported in the 2008 nationwide survey (11). According to the calculation, increased blood pressure (> 130/85 mmHg) yielded the sample size estimate of 386 at 5% level of significance and a precision of 0.05.

Frequencies and percentages were obtained for categorical variables. Chi square tests for Independence were used to determine the univariate association between categorical variables. Multiple logistic regression analyses was used to determine the associations of sociodemographic and lifestyle factors with MetS, calculating odds ratios with 95% confidence intervals, while adjusting for confounding factors. Variables, with p < 0.25 on univariate analysis were selected for adjustment in the final logistic regression model. P < 0.05 was considered statistically significant.

### Ethics

Ethical approval was sought from the Monash University Human Research Ethics Committee (Project # CF15/56-2016000022).

## Results

The prevalence of MetS was found to be 32.2% in the study subjects, according to the Harmonized criteria; highest among the Indians (51.9%) and lowest among the Chinese (20.2%) (Fig 1). Abdominal obesity (62.0%) and high blood pressure (56.8%) were more common compared to other metabolic abnormalities. Three most prominent MetS risk factors among Malays and Indians were abdominal obesity, high blood pressure and low HDL-cholesterol. Among the Indians however, the percentages of abdominal obesity and HDL-cholesterol were higher than that among the Malays. Prevalence of high blood pressure was more prominent among the Malays compared to the other ethnic groups. Among the Chinese, the third most prevalent risk factor was hypertriglyceridemia. Prevalence of low HDL-cholesterol was at its lowest among the Chinese. Table 1 shows the comparative association of sociodemographic and lifestyle characteristics with MetS overall, and across the 3 major ethnicities in Johor. Overall, significant differences are observed with age, ethnicity, marital status, education and physical activity (p < 0.05). Marital status and education were found to be significantly related with MetS among the Malays, while age and physical activity among the Chinese and age among the Indians showed a significant association with MetS. Among the Malays, 59.4% of the people with primary education or lower were having MetS, suggesting that those Malays having higher education appear to be protected against the risk of MetS (p < 0.05). Majority of the Indians appear to be afflicted with MetS at a younger age (41.6% at the age group of 40-49 years; p < 0.001). Conversely, only 15.2% of the Chinese were suffering from this syndrome in this age group (p = 0.016). This shows that the Chinese in Johor are getting this disease at a relatively older age.

**Fig 1:**
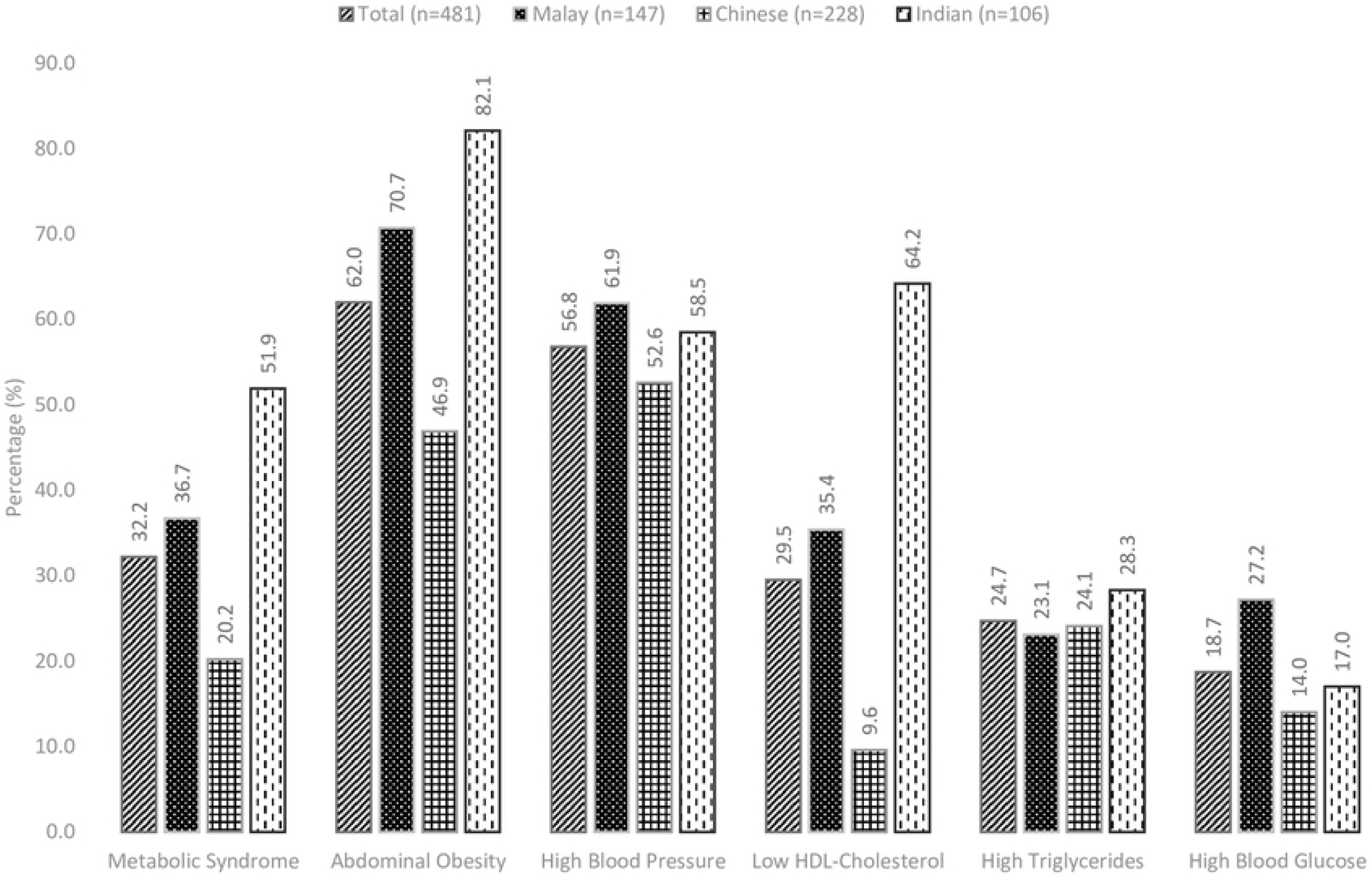
Proportions of MetS and its components, overall and among the three major ethnic groups in Johor

**Table 1:** Summary of sociodemographic, lifestyle and dietary characteristics with MetS, overall and among the three major ethnic groups in Johor

|  |  | Overall |  |  | Malay |  |  | Chinese |  |  | Indian |  |  |
| --- | --- | --- | --- | --- | --- | --- | --- | --- | --- | --- | --- | --- | --- |
|  |  | Total<br>(n=481) | With MetS<br>(n=155) | P-value* | Total<br>(n=147) | With MetS<br>(n=54) | P-value* | Total<br>(n=228) | With MetS<br>(n=46) | P-value* | Total<br>(n=106) | With MetS<br>(n=55) | P-value* |
| Characteristic |  | n (%) | n (%) |  | n (%) | n (%) |  | n (%) | n (%) |  | n (%) | n (%) |  |
| Gender | Male | 169 (35.1) | 57 (36.8) | 0.604 | 42 (28.6) | 12 (22.2) | 0.194 | 88 (38.6) | 20 (43.5) | 0.447 | 39 (36.8) | 25 (45.5) | 0.055 |
|  | Female | 312 (64.9) | 98 (63.2) |  | 105 (71.4) | 42 (77.8) |  | 140 (61.4) | 26 (56.5) |  | 67 (63.2) | 30 (54.5) |  |
| Age (years) | < 40 | 112 (23.3) | 22 (14.2) | 0.005 | 38 (25.9) | 12 (22.2) | 0.442 | 40 (17.5) | 3 (6.5) | 0.016 | 34 (32.1) | 7 (12.7) | < 0.001 |
|  | 40-49 | 129 (26.8) | 48 (31.0) |  | 56 (38.1) | 18 (33.3) |  | 35 (15.4) | 7 (15.2) |  | 38 (35.8) | 23 (41.8) |  |
|  | 50-59 | 117 (24.3) | 36 (23.2) |  | 32 (21.8) | 14 (25.9) |  | 71 (31.1) | 11 (23.9) |  | 14 (13.2) | 11 (20.0) |  |
|  | ≥ 60 | 123 (25.6) | 49 (31.6) |  | 21 (14.3) | 10 (18.5) |  | 82 (36.0) | 25 (54.3) |  | 20 (18.9) | 14 (25.5) |  |
| Ethnicity | Malay | 147 (30.6) | 54 (34.8) | < 0.001 | - | - | - | - | - | - | - | - | - |
|  | Chinese | 228 (47.4) | 46 (29.7) |  | - | - |  | - | - |  | - | - |  |
|  | Indian | 106 (22.0) | 55 (35.5) |  | - | - |  | - | - |  | - | - |  |
| Marital status | Single | 63 (13.1) | 13 (8.4) | 0.002 | 5 (3.4) | 3 (5.6) | 0.035 | 49 (21.5) | 7 (15.2) | 0.284 | 9 (8.5) | 3 (5.5) | 0.420 |
|  | Married/living with partner | 378 (78.6) | 123 (79.4) |  | 126 (85.7) | 41 (75.9) |  | 168 (73.7) | 38 (82.6) |  | 84 (79.2) | 44 (80.0) |  |
|  | Separated/divorced | 40 (8.3) | 19 (12.3) |  | 16 (10.9) | 10 (18.5) |  | 11 (4.8) | 1 (2.2) |  | 13 (12.3) | 8 (14.5) |  |
| Education | Primary or lower | 95 (19.8) | 47 (30.3) | < 0.001 | 32 (21.8) | 19 (35.2) | 0.004 | 35 (15.4) | 12 (26.1) | 0.062 | 28 (26.4) | 16 (29.1) | 0.255 |
|  | Secondary | 291 (60.5) | 92 (59.4) |  | 100 (68.0) | 33 (61.1) |  | 126 (55.3) | 24 (52.2) |  | 65 (61.3) | 35 (63.6) |  |
|  | Tertiary | 95 (19.8) | 16 (10.3) |  | 15 (10.2) | 2 (3.7) |  | 67 (29.4) | 10 (21.7) |  | 13 (12.3) | 4 (7.3) |  |
| Employment status | Employed | 206 (42.8) | 61 (39.4) | 0.289 | 47 (32.0) | 14 (25.9) | 0.231 | 111 (48.7) | 20 (43.5) | 0.429 | 48 (45.3) | 27 (49.1) | 0.413 |
|  | Unemployed | 275 (57.2) | 94 (60.6) |  | 100 (68.0) | 40 (74.1) |  | 117 (51.3) | 26 (56.5) |  | 58 (54.7) | 28 (50.9) |  |
| Physical activity | High | 95 (19.8) | 26 (16.8) | 0.045 | 66 (44.9) | 27 (50.0) | 0.520 | 81 (35.5) | 23 (50.0) | 0.029 | 47 (44.3) | 25 (45.5) | 0.957 |
|  | Moderate | 192 (39.9) | 54 (34.8) |  | 52 (35.4) | 16 (29.6) |  | 102 (44.7) | 19 (41.3) |  | 38 (35.8) | 19 (34.5) |  |
|  | Low | 194 (40.3) | 75 (48.4) |  | 29 (19.7) | 11 (20.4) |  | 45 (19.7) | 4 (8.7) |  | 21 (19.8) | 11 (20.0) |  |
| Smoking status | Never smoked | 419 (87.1) | 138 (89.0) | 0.386 | 124 (84.4) | 49 (90.7) | 0.104 | 205 (89.9) | 43 (93.5) | 0.369 | 90 (84.9) | 46 (83.6) | 0.705 |
|  | Past/Current smoker | 62 (12.9) | 17 (11.0) |  | 23 (15.6) | 5 (9.3) |  | 23 (10.1) | 3 (6.5) |  | 16 (15.1) | 9 (16.4) |  |
| Alcohol consumption | Never/past consumer | 440 (91.5) | 139 (89.7) | 0.330 | 146 (99.3) | 49 (90.7) | 0.104 | 205 (89.9) | 43 (93.5) | 0.369 | 90 (84.9) | 46 (83.6) | 0.705 |
|  | Current consumer | 41 (8.5) | 16 (10.3) |  | 1 (0.7) | 1 (1.9) |  | 23 (10.1) | 3 (6.5) |  | 16 (15.1) | 9 (16.4) |  |
| Frequent dining out | ≤ 3 times/week | 327 (68.0) | 110 (71.0) | 0.333 | 132 (89.8) | 47 (87.0) | 0.400 | 110 (48.2) | 22 (47.8) | 0.949 | 85 (80.2) | 41 (74.5) | 0.130 |
|  | > 3 times/week | 154 (32.0) | 45 (29.0) |  | 15 (10.2) | 7 (13.0) |  | 118 (51.8) | 24 (52.2) |  | 21 (19.8) | 14 (25.5) |  |
| Late dining | ≤ 3 times/week | 428 (89.0) | 136 (87.7) | 0.549 | 129 (87.8) | 49 (90.7) | 0.400 | 208 (91.2) | 41 (89.1) | 0.574 | 91 (85.8) | 46 (83.6) | 0.497 |
|  | **> 3 times/week** | 53 (11.0) | 19 (12.3) |  | 18 (12.2) | 5 (9.3) |  | 20 (8.8) | 5 (10.9) |  | 15 (14.2) | 9 (16.4) |  |
| **Skipping breakfast** | **≤ 3 times/week** | 364 (75.7) | 119 (76.8) | 0.699 | 117 (79.6) | 42 (77.8) | 0.678 | 175 (76.8) | 37 (80.4) | 0.508 | 72 (67.9) | 40 (72.7) | 0.271 |
|  | **> 3 times/week** | 117 (24.3) | 36 (23.2) |  | 30 (20.4) | 12 (22.2) |  | 53 (23.2) | 9 (19.6) |  | 34 (32.1) | 15 (27.3) |  |
| **Finishing meals** | **Not fast** | 229 (47.6) | 66 (42.6) | 0.128 | 79 (53.7) | 24 (44.4) | 0.085 | 121 (53.1) | 24 (52.2) | 0.892 | 29 (27.4) | 18 (32.7) | 0.198 |
|  | **Fast** | 252 (52.4) | 89 (57.4) |  | 68 (46.3) | 30 (55.6) |  | 107 (46.9) | 22 (47.8) |  | 77 (72.6) | 37 (67.3) |  |

\*P-value ascertained by $\chi^2$ test

Table 2 shows the adjusted multiple logistic regression model, the results of which indicate that overall in this population, higher age groups and the Indian race had increased the odds of having MetS, while the Chinese ethnicity and tertiary education were protective against the risk of MetS. Lifetsyle factors and the diet habits did not appear to have any association with MetS, overall, in the adjusted model (p > 0.05).

**Table 2:** Association of sociodemographic and lifestyle factors and diet habits with MetS among the overall population of Johor (n=481)

|  | Characteristic | Crude OR (95% CI) | Adjusted OR (95% CI) |
| --- | --- | --- | --- |
| **Gender** | **Male** | 1.00 |  |
|  | **Female** | 0.90 (0.60-1.34) |  |
| **Age (years)** | **< 40** | 1.00 | 1.00 |
|  | **40-49** | 2.42 (1.35-4.36)* | 2.44 (1.27-4.70)* |
|  | **50-59** | 1.82 (0.99-3.34) | 2.80 (1.36-5.79)* |
|  | **≥ 60** | 2.71 (1.50-4.88)* | 4.14 (1.97-8.69)* |
| **Ethnicity** | **Malay** | 1.00 | 1.00 |
|  | **Chinese** | 0.44 (0.27-0.69)* | 0.37 (0.21-0.65)* |
|  | **Indian** | 1.86 (1.12-3.08)* | 1.95 (1.12-3.38)* |
| **Marital status** | **Single** | 1.00 | 1.00 |
|  | **Married/living with partner** | 1.86 (0.97-3.54) | 0.81 (0.38-1.71) |
|  | **Separated/divorced** | 3.48 (1.46-8.31)* | 0.75 (0.26-2.19) |
| **Education** | **Primary or lower** | 1.00 | 1.00 |
|  | **Secondary** | 0.47 (0.29-0.76)* | 0.63 (0.37-1.08) |
|  | **Tertiary** | 0.21 (0.11-0.40)* | 0.44 (0.20-0.94)* |
| **Employment status** | **Employed** | 1.00 |  |
|  | **Unemployed** | 1.23 (0.84-1.82) |  |
| **Physical activity** | **High** | 1.00 | 1.00 |
|  | **Moderate** | 1.67 (0.98-2.86) | 1.14 (0.63-2.04) |
|  | **Low** | 1.04 (0.60-1.80) | 1.77 (0.99-3.16) |
| **Smoking status** | **Never smoked** | 1.00 |  |
|  | **Past/Current smoker** | 0.77 (0.42-1.39) |  |
| **Alcohol consumption** | **Never consumed/past consumer** | 1.00 |  |
|  | **Current consumer** | 1.39 (0.72-2.68) |  |
| **Frequent dining out** | **≤ 3 times/week** | 1.00 |  |
|  | **> 3 times/week** | 0.81 (0.54-1.24) |  |
| **Late dining** | **≤ 3 times/week** | 1.00 |  |
|  | **> 3 times/week** | 1.20 (0.66-2.18) |  |
| **Skipping breakfast** | **≤ 3 times/week** | 1.00 |  |
|  | **> 3 times/week** | 0.92 (0.58-1.43) |  |
| **Finishing meals** | **Not fast** | 1.00 | 1.00 |
|  | **Fast** | 1.35 (0.92-1.98) | 1.15 (0.75-1.78) |

\*P-value < 0.05
OR=Odds ratio; CI=Confidence interval; Empty cells refers to the variables excluded ( $p > 0.25$ on univariate analyses) from the model

In view of the contrasting estimates of MetS among the ethnicities, we explored the effect of ethnicity further with sociodemographic, lifestyle and diet factors. Table 3 shows the adjusted logistic regression models among the Malays, the Chinese and the Indians, revealing higher odds for MetS for quick finishing of meals among the Malays (AOR = 2.17 [95% CI = 1.02-4.60]) and low physical activity among the Chinese (AOR = 4.76 [95% CI = 1.49-15.26]). Furthermore, higher educational categories were protective against MetS among the Malays. Among the Indians, older age groups (40 years and above) were more prone to developing MetS, while significant odds with respect to age were found among the Chinese older than 60 years of age (AOR = 5.43 [95% CI = 1.39 – 21.13]).

**Table 3:**
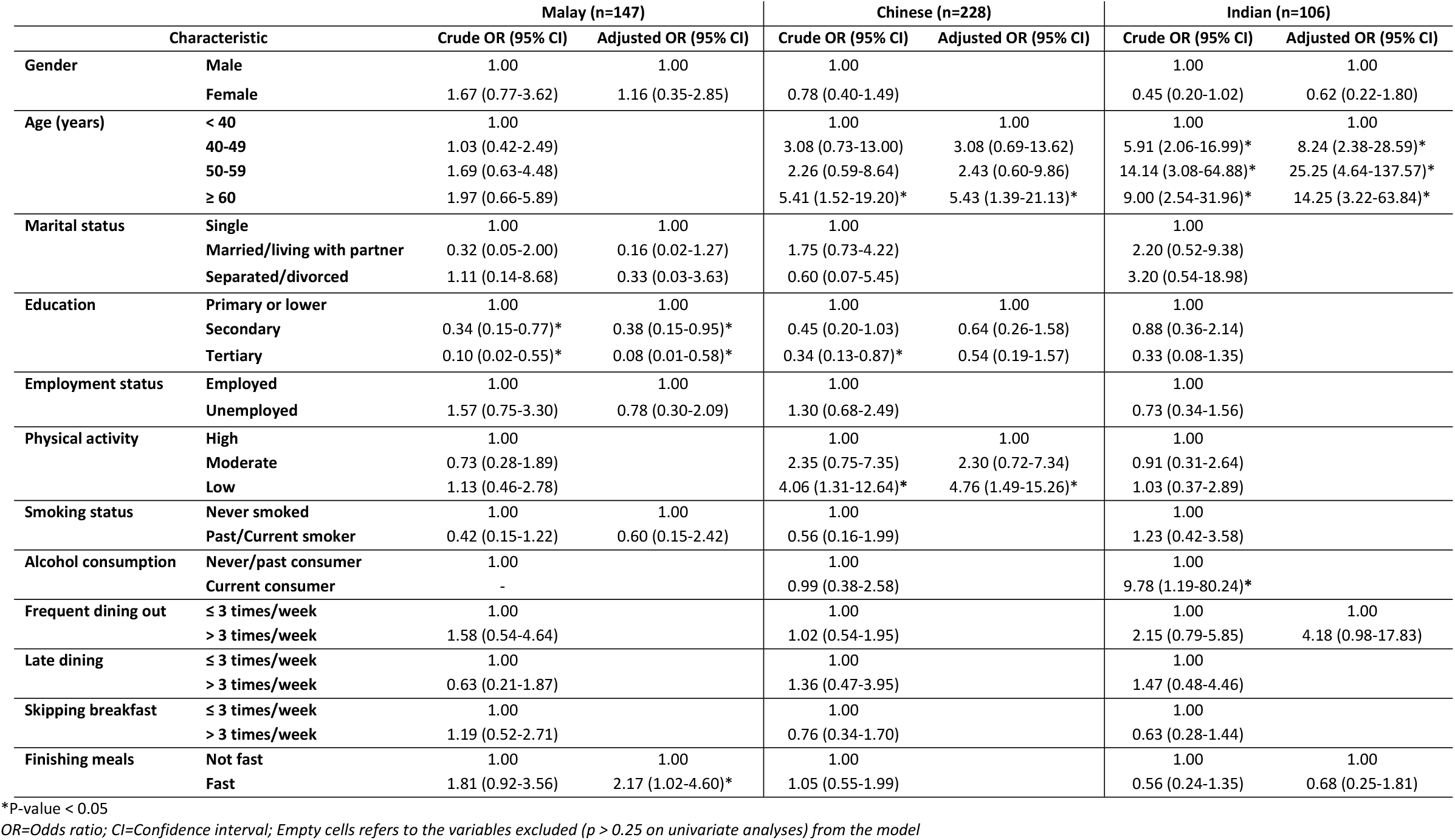
Association of sociodemographic and lifestyle factors and diet habits with MetS among the three major ethnicities of Johor

|  |  | Malay (n=147) |  | Chinese (n=228) |  | Indian (n=106) |  |
| --- | --- | --- | --- | --- | --- | --- | --- |
| Characteristic |  | Crude OR (95% CI) | Adjusted OR (95% CI) | Crude OR (95% CI) | Adjusted OR (95% CI) | Crude OR (95% CI) | Adjusted OR (95% CI) |
| Gender | Male | 1.00 | 1.00 | 1.00 |  | 1.00 | 1.00 |
|  | Female | 1.67 (0.77-3.62) | 1.16 (0.35-2.85) | 0.78 (0.40-1.49) |  | 0.45 (0.20-1.02) | 0.62 (0.22-1.80) |
| Age (years) | < 40 | 1.00 |  | 1.00 | 1.00 | 1.00 | 1.00 |
|  | 40-49 | 1.03 (0.42-2.49) |  | 3.08 (0.73-13.00) | 3.08 (0.69-13.62) | 5.91 (2.06-16.99)* | 8.24 (2.38-28.59)* |
|  | 50-59 | 1.69 (0.63-4.48) |  | 2.26 (0.59-8.64) | 2.43 (0.60-9.86) | 14.14 (3.08-64.88)* | 25.25 (4.64-137.57)* |
|  | ≥ 60 | 1.97 (0.66-5.89) |  | 5.41 (1.52-19.20)* | 5.43 (1.39-21.13)* | 9.00 (2.54-31.96)* | 14.25 (3.22-63.84)* |
| Marital status | Single | 1.00 | 1.00 | 1.00 |  | 1.00 |  |
|  | Married/living with partner | 0.32 (0.05-2.00) | 0.16 (0.02-1.27) | 1.75 (0.73-4.22) |  | 2.20 (0.52-9.38) |  |
|  | Separated/divorced | 1.11 (0.14-8.68) | 0.33 (0.03-3.63) | 0.60 (0.07-5.45) |  | 3.20 (0.54-18.98) |  |
| Education | Primary or lower | 1.00 | 1.00 | 1.00 | 1.00 | 1.00 |  |
|  | Secondary | 0.34 (0.15-0.77)* | 0.38 (0.15-0.95)* | 0.45 (0.20-1.03) | 0.64 (0.26-1.58) | 0.88 (0.36-2.14) |  |
|  | Tertiary | 0.10 (0.02-0.55)* | 0.08 (0.01-0.58)* | 0.34 (0.13-0.87)* | 0.54 (0.19-1.57) | 0.33 (0.08-1.35) |  |
| Employment status | Employed | 1.00 | 1.00 | 1.00 |  | 1.00 |  |
|  | Unemployed | 1.57 (0.75-3.30) | 0.78 (0.30-2.09) | 1.30 (0.68-2.49) |  | 0.73 (0.34-1.56) |  |
| Physical activity | High | 1.00 |  | 1.00 | 1.00 | 1.00 |  |
|  | Moderate | 0.73 (0.28-1.89) |  | 2.35 (0.75-7.35) | 2.30 (0.72-7.34) | 0.91 (0.31-2.64) |  |
|  | Low | 1.13 (0.46-2.78) |  | 4.06 (1.31-12.64)* | 4.76 (1.49-15.26)* | 1.03 (0.37-2.89) |  |
| Smoking status | Never smoked | 1.00 | 1.00 | 1.00 |  | 1.00 |  |
|  | Past/Current smoker | 0.42 (0.15-1.22) | 0.60 (0.15-2.42) | 0.56 (0.16-1.99) |  | 1.23 (0.42-3.58) |  |
| Alcohol consumption | Never/past consumer | 1.00 |  | 1.00 |  | 1.00 |  |
|  | Current consumer | - |  | 0.99 (0.38-2.58) |  | 9.78 (1.19-80.24)* |  |
| Frequent dining out | ≤ 3 times/week | 1.00 |  | 1.00 |  | 1.00 | 1.00 |
|  | > 3 times/week | 1.58 (0.54-4.64) |  | 1.02 (0.54-1.95) |  | 2.15 (0.79-5.85) | 4.18 (0.98-17.83) |
| Late dining | ≤ 3 times/week | 1.00 |  | 1.00 |  | 1.00 |  |
|  | > 3 times/week | 0.63 (0.21-1.87) |  | 1.36 (0.47-3.95) |  | 1.47 (0.48-4.46) |  |
| Skipping breakfast | ≤ 3 times/week | 1.00 |  | 1.00 |  | 1.00 |  |
|  | > 3 times/week | 1.19 (0.52-2.71) |  | 0.76 (0.34-1.70) |  | 0.63 (0.28-1.44) |  |
| Finishing meals | Not fast | 1.00 | 1.00 | 1.00 |  | 1.00 | 1.00 |
|  | Fast | 1.81 (0.92-3.56) | 2.17 (1.02-4.60)* | 1.05 (0.55-1.99) |  | 0.56 (0.24-1.35) | 0.68 (0.25-1.81) |

\*P-value < 0.05
OR=Odds ratio; CI=Confidence interval; Empty cells refers to the variables excluded ( $p > 0.25$ on univariate analyses) from the model

## Discussion

In this study, the prevalence of MetS was found to be 32.2%, which was unexpectedly less than that reported in the 2008 nationwide survey, that contained 19% of subjects from Johor (11). Prevalence among the Indians and the Chinese turned out to be 51.9% and 20.2%, respectively. Comparing the prevalence values reported in the nationwide survey 2008 for these 2 ethnic groups in Malaysia, the prevalence among the Indians appears to have remained unchanged over a period of 9 years, however among the Chinese, the prevalence has reduced considerably from 42.1% to 20.2%, while among the Malays the prevalence has decreased from 43.9% to 36.7% (11). This apparent decline among the Malays and the Chinese may be attributed to the decreased prevalence values of hyperglycemia, low HDL-cholesterol and hypertriglyceridemia.

The present study showed that MetS was more prominent among the higher age groups. This finding has been observed by other researchers also. He et al., reported a comparatively higher prevalence of MetS among older subjects (70 years and above) compared to those aged between 60-69 years among a total of over two thousand Chinese subjects. (38). In the study by Rampal et al., the prevalence of MetS among Malaysians was found to be higher among subjects aged 40 years and above compared to subjects less than 40 years (44.6% vs. 16.0%) (39). Also, in the study by Ramli and colleagues, the odds of MetS, irrespective of definition applied, were found to be higher among higher age groups, and maximum among subjects aged 60 years and above (12). The nationwide survey also reported higher prevalence of MetS among higher age groups; additionally, higher age groups also had a higher prevalence of central obesity, high blood pressure, low HDL-cholesterol, elevated triglycerides and hyperglycemia (11). This suggests that higher prevalence of MetS among higher age groups may be due to the accumulated higher prevalence of its associated cardio-metabolic risk factors among elderly subjects.

Studies have shown that ethnicity influences (directly or indirectly) the prevalence of metabolic syndrome. For instance, from a study in Canada, MetS prevalence was reported to be higher among the Cree Indians compared to other aboriginal and non-aboriginal Canadians (14). The Cree Indians also had a higher prevalence of central obesity and hyperglycemia compared to other races in the country (14). Similarly, from a study in Suriname, South America, MetS prevalence was reported as the highest among the Hindustanis (descendent of Indians), compared to other Suriname races (40). The prevalence values of high blood pressure, low HDL-cholesterol and hyperglycemia were also high among the Suriname Hindustanis (40). In our study, results show that the Chinese appear to be less prone to developing MetS, while the Indians in Johor are at a greater risk of developing MetS. This is in line with the reports from other researchers from Malaysia that the Chinese have lower odds, while the Indians have higher odds of developing MetS (12, 39). More educated adults in the Johor area, especially Malays, appear to be protected against MetS, probably due to their more awareness of healthy lifestyle habits, such as physical activity, smoking cessation, moderate to none consumption of alcohol and adoption of healthy eating habits (32). This is supported by a couple of studies showing that higher education of individuals is protective against diabetes and hypertension, which are prominent risk components of MetS (41, 42). However, Ching et al., have recently reported that higher education levels of Malaysian vegetarians with and without MetS were nearly the same (43). This could be due to the fact that that was a unique group of subjects with specific dietary habits and the results pertaining to this group may not represent the general population of Malaysia.

Literature suggests that excess energy accumulated in the adipose tissues causes metabolic abnormalities, leading to high blood pressure, hyperglycemia, hypertriglyceridemia and inflammation, hence, regular physical activity enhances energy consumption leading to a reduced prevalence of obesity, hypertension, diabetes mellitus and also MetS (44, 45). Our results show the prevalence of MetS and its components to be comparatively lower among Chinese than in Malays and Indians, which may be likely attributed to better lifestyle choices, including physical activity. Chu et al., have shown that longer sitting time and insufficient physical activity have resulted in an almost 4-fold increase in MetS risk among the Malays, and the risk gets reduced by 50% by engaging in moderate to high physical activity (27, 28). On the basis of these reports, it can be suggested that the Chinese in Johor, though having a decreased risk of developing MetS, can still benefit by engaging in moderate to high levels of physical activity.

Although a number of studies have shown a direct relationship of smoking with the risk of MetS, yet in the current study smoking does not appear to be associated with the risk of MetS (20). This could be due to a small proportion of past and current smokers (12.9%) in this cohort. Similarly, no association was found between alcohol consumption and risk of MetS in this population. Again the reason could be the small proportion of subjects who reported as alcohol consumers (8.5%). The association of dietary habits, such as quick finishing of meals, frequent dining out, late eating, skipping breakfast, with MetS has been reported in other studies in the East Asian region (25, 26, 46–49). For example, Shin et al., reported faster eating as one of the risk factors for MetS among the Koreans (49). Among these dietary habits, quick finishing of meals was identified in the current study as a new risk factor for MetS in Malaysia, especially among the Malays. According to Dallman and colleagues, fast eaters may consume more food than usual, or be eating under psychological stress which affects hormones regulating metabolism (50). The underlying mechanism of such habit(s) with the metabolic health functioning, however remains unclear (49). Certain limitations warrant consideration. First, the present research study was cross-sectional in nature, assessing the exposures and outcomes at the same point in time. In this regard, the findings cannot indicate causality. Second, the selection of study locations harboring subjects was nonrandom and partly based on the available information on the percentage of MetS across each Malaysian ethnicity in Johor reported in the nationwide survey 2008 (11). Where this was done to have sufficient numbers of subjects in each ethnic group for better analysis and interpretation, however, a selection bias cannot be completely discounted. Furthermore, as the information collected was based on recall, hence misreporting of information cannot be completely ruled out, and this might have added some variability in our results. The variable effects are more likely attributed to the cultural diversities across the different ethnicities of Malaysia. Despite these sources of potential variability, the results provide evidence towards the association of certain sociodemographic, lifestyle and diet factors that affect the disease spectrum of MetS in Johor and with a reasonable sample size, it did provide an opportunity to have an in-depth analysis of sociodemographic and lifestyle factors and diet habits influencing MetS across the three major races of this state in the country. In this regard, we believe the research to be adequate and its findings comparable to similar studies by other investigators using a non-randomized design exploring associations of various risk factors influencing metabolic diseases within Malaysia and abroad.

Population in Johor is diverse in its habits pertaining to lifestyle and diet. Some of these factors are associated with the risk of MetS in certain ethnic groups, and modifying these factors would be important for reducing cardiovascular and metabolic health risks in the country. Malaysia is a multi-ethnic country, and it would be important to consider the ethnic variation, especially with respect to lifestyle and diet factors, hence, intervention programs would require to be tailored across the different races of the country for addressing behavior modifications. Increasing awareness among the masses through electronic and print media about the beneficial effects of healthy lifestyle is likely to be a very powerful approach to combat the menace of this syndrome in this country. Moreover, further prospective studies delineating the association of various diet habits among different racial communities are needed to contain the unfavorable effects of this syndrome on the overall health of Malaysians.

## Acknowledgements

Financial assistance was provided by the Clinical Research Center, Monash University, Malaysia, and by the grant awarded to Dr Amutha Ramadas by the Malaysian Ministry of Higher Education’s Fundamental Research Grant Scheme (FRGS/2/2013/SKK07/MUSM/03/1). Also, the authors wish to thank Gribbles Laboratories (MS ISO 15189), Malaysia, for assisting with the laboratory assessments on the collected blood samples. We also appreciate all the assistance from Government contacts (Majlis Perbandaran Kulai) and all local community leaders. We would also like to extend our gratitude to the Monash University faculty and staff, namely Mr Chui Chor Sin, Mrs Savithri Gopal, Ms Pang Pei Ling, Ms Ungku Zulaikha Ungku Omar, Mr Muhammad Daniel Mahadzir, Dr Nor Azim and Dr Iekhsan Othman for all their support and assistance in this study.

## Authors’ contributions

Study conceptualization: SPI, AR, QKF and KAK.

Population health screening: HLS and WYO

Project administration: SPI, AR, HLS, WYO and KAK

Data analysis and interpretation: SPI and KAK.

Manuscript writing and review: SPI, AR, QKF and KAK.

